# Photothermally Detected Stimulated Raman Microscopy towards Ultrasensitive Chemical Imaging

**DOI:** 10.1101/2023.03.06.531387

**Authors:** Yifan Zhu, Xiaowei Ge, Hongli Ni, Jiaze Yin, Haonan Lin, Le Wang, Yuying Tan, Chinmayee V. Prabhu Dessai, Ji-Xin Cheng

## Abstract

Stimulated Raman scattering (SRS) microscopy has shown enormous potential in revealing molecular structures, dynamics and couplings in complex systems. However, the sensitivity of SRS is fundamentally limited to milli-molar level due to the shot noise and the small modulation depth. To overcome this barrier, we revisit SRS from the perspective of energy deposition. The SRS process pumps molecules to their vibrationally excited states. The thereafter relaxation heats up the surrounding and induces refractive index changes. By probing the refractive index changes with a laser beam, we introduce stimulated Raman photothermal (SRP) microscopy, where a >500-fold boost of modulation depth is achieved. Versatile applications of SRP microscopy on viral particles, cells, and tissues are demonstrated. SRP microscopy opens a new way to perform vibrational spectroscopic imaging with ultrahigh sensitivity.

**One-Sentence Summary:** We demonstrate a new spectroscopic imaging method that improves the signal intensity by >500-fold.

## Main Text

Avoiding the non-resonant background encountered in coherent anti-Stokes Raman scattering (CARS) microscopy, stimulated Raman scattering (SRS) microscopy has become a powerful platform in the field of chemical imaging (*1–3*). In SRS (Fig. 1A), two spatial-temporally overlapped laser pulse trains, namely pump and Stokes, coherently interact with Raman-active molecules that resonates at the laser beating frequency, causing a stimulated Raman gain and loss in the Stokes and the pump field, respectively. While providing the same molecular vibrational features to conventional Raman spectroscopy, SRS offers up to 6 orders of speed improvement comparing to spontaneous Raman (*3*). Such speed improvement, together with development of vibrational probes (*4*), has enabled many applications, e.g. label-free stimulated Raman histology (*5,6*), nutrient mapping (*7–9*), unveiling altered metabolism in cancer (*10–14*), heterogeneity in microbiome (*15*), rapid detection of antimicrobial susceptibility (*16*). Despite these advances, the detection sensitivity of SRS is fundamentally challenged by the small modulation depth (0.1% for a pure liquid) and the shot noise in the pump or Stokes beam (*17,18*), where simply increasing the number of photons can easily exceed the power tolerance of the sample.

**Fig. 1.**
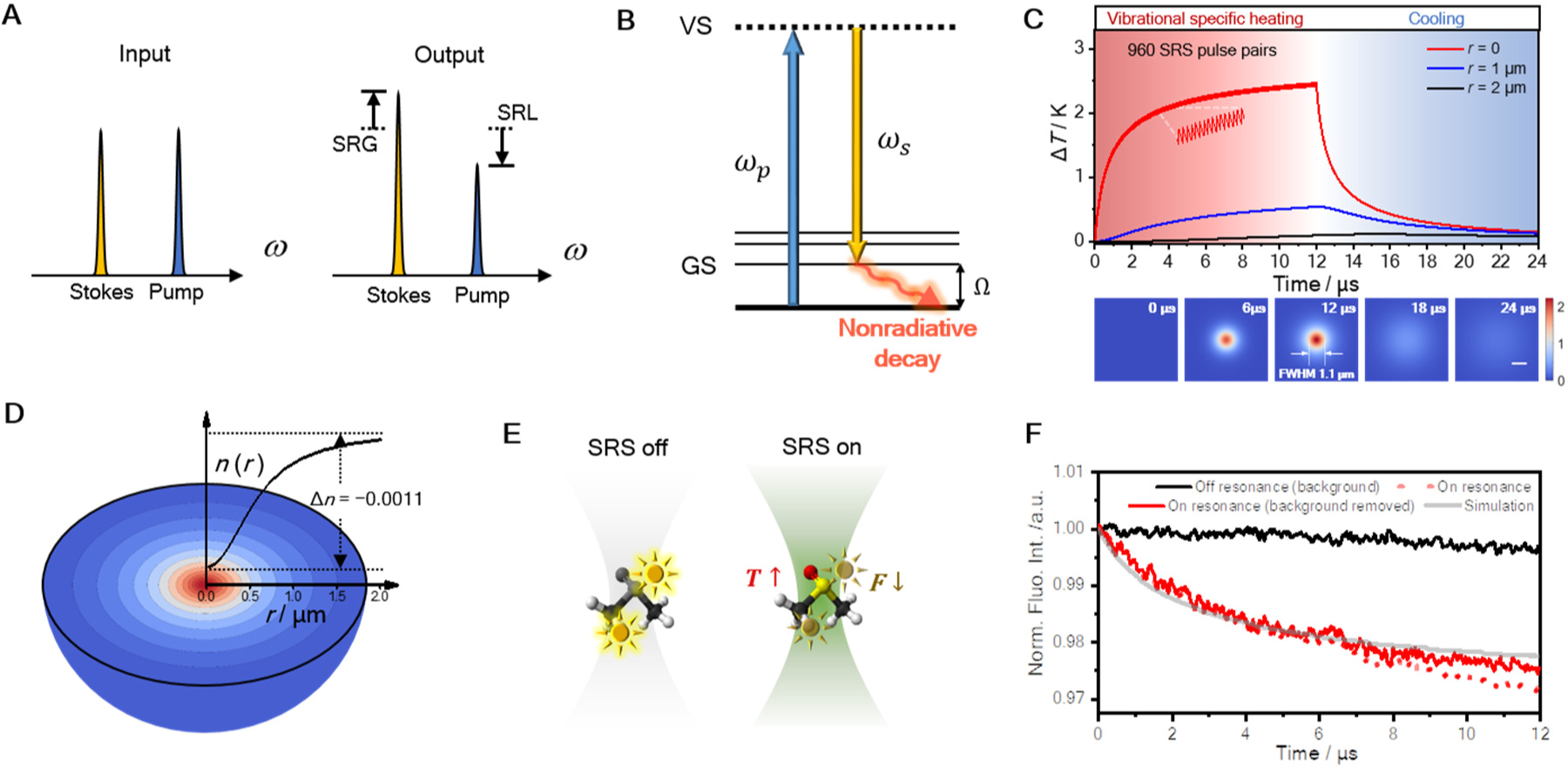
Theoretical simulation and experimental observation of the SRP effect. (**A**) Schematic of stimulated Raman gain and loss. (**B**) Schematic of stimulated Raman photothermal (SRP) effect. (**C**) Simulation of temperature rise induced by SRP in temporal (top) and spatial (bottom) domains. Spatial scale bar: 1 μm. (**D**) Simulated profile of thermal lens induced by SRP in pure DMSO. (**E**) Illustration of fluorescence thermometer measurement of SRP-mediated temperature rise. (**F**) Measured fluorescence of rhodamine B in DMSO during an SRS process. The beating frequency (*ω*_p_−*ω*_s_) is tuned to 2913cm^−1^ for on-resonance and 2850 cm^−1^ for off-resonance.

Pushing the fundamental limit of SRS sensitivity requires either a reduction of measurement noise, or a boost of SRS signal. On the reduction of SRS measurement noise, efforts focused on using squeezed light, termed “quantum-enhanced SRS”. Signal-to-noise-ratio (SNR) enhancements of 3.6 dB (*19*) with continuous wave squeezed light, or 2.89 dB (*20*) with pulsed squeezed light have been demonstrated with no additional perturbation on sample. While the future of this method is bright, it is currently limited by the low squeeze efficiency and decoherence in complex imaging systems. On the boost of SRS signal, different photophysical processes have been utilized to increase the cross-section, hence the signal intensity, including electronic pre-resonance SRS (*21–23*), plasmon enhanced SRS (*24*) and stimulated Raman excited fluorescence (*25*). Very high enhancement factor (10^4^ ∼ 10^7^) of SRS signal, and single molecule SRS measurement (*24,25*), have been achieved. However, the requirement of special target molecules or plasmonic nanostructures constrains the scope of applications.

To seek new approaches towards boosting the signal, we revisit the physics of SRS from the perspective of energy transfer from laser fields to the sample. As explored by Cheng and coworkers in 2007 from the perspective of photodamage in CARS imaging (*26*), ∼0.08% of the laser power is absorbed by the sample through the simultaneously occurred stimulated Raman gain and loss processes. As illustrated in Fig. 1**b**, when pump and Stokes pulses with appropriate wavelengths interact with Raman-active molecules, the target molecules are pumped to their vibrationally excited states, with the transition energy equals to the beating frequency between the pump and Stokes lasers. Importantly, after SRS excitation, the vibrationally excited molecules relax their vibrational energy quickly through non-radiative decay, and consequently, heat up the surrounding environment, causing a stimulated Raman photothermal (SRP) effect.

Optically detected photothermal microscopy has been well developed (*27*) and has reached the sensitivity down to a single molecule (*28*). In photothermal spectroscopy, first reported in 1970s (*29*), an optical absorption raises the local temperature, creating local change of refractive index, which is measured with a probing beam. Early photothermal microscopy works focused on electronic absorption, targeting non-fluorescent dye molecules or metal nanostructures (*27*).

Recently developed mid-infrared photothermal (MIP) microscopy provides universal infrared active vibrational spectroscopic features, offers a detection sensitivity at micro-molar level, and creates a spatial resolution at the visible diffraction limit (*30,31*) or even higher by probing the high harmonic signals (*32*). To the contrary, the thermal effects induced by a Raman process is commonly believed minimal due to the small cross sections of Raman scattering, and consequent Raman based photothermal imaging is usually considered impossible. However, with coherent Raman excitation that nonlinearly benefits from high laser peak power, we prove and quantify the thermal effect of SRS process, and demonstrate its potential in chemical imaging with ultrahigh sensitivity, complementary spectroscopic information to MIP, as well as negligible water absorption.

## Results and discussion

### Simulation of the SRP effect

For a stimulated Raman loss measurement, the relationship between pump beam modulation depth η, pump laser intensity *I*_p_, and change of photon number per pulse, *ΔN*, can be written as:

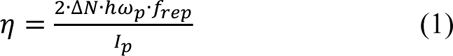

where *h* is the Plank’s constant, *ω*_p_ is the pump wavenumber, and *f*_rep_ is the laser repetition rate. With this, one can estimate the energy deposition per pair of SRS pulses by *E* = Δ*N* · ℎ*ω*_R_, where *ω*_R_ is the target Raman shift. Literature (*4*) has shown that with 25 mW (modulated at 50% duty cycle) and 15 mW on-sample powers for 80 MHz Stokes and pump beams, respectively, the SRS modulation depth on the 2913 cm^-1^ mode of dimethyl sulfoxide (DMSO) reaches 0.04%. By inserting these measured values into the equations, the energy deposition per pair of laser pulses is found to be 8.7 fJ, equivalent to 0.7 μW.

With this energy deposition estimation, we applied Fourier’s law and built a finite unit model (Fig. S1) to quantitatively simulate the stimulated Raman induced temperature rise in pure DMSO. Simulation results (Fig. 1C) show that, when using an objective with 0.8 N.A. (numerical aperture), and routinely used laser power (25 mW and 50 mW for pump and Stokes, 80 MHz repetition rate, 50% duty cycle), the temperature rise at the center of the laser focus can reach as high as 2.4 K after 12 µs of stimulated Raman excitation, corresponding to 960 pairs of pump/Stokes pulses. The temperature rises at r = 1 μm and 2 μm away from the focal center are 0.54 K and 0.12 K, respectively, at t = 12 μs. The temperature map at different time points is also shown in Fig. 1C. The full width at half maximum (FWHM) of the temperature rising field at t = 12 μs is calculated to be 1.1 μm, suggesting a very localized thermal field.

The temperature elevation subsequently changes the local refractive index through the thermal-optic effect. For pure DMSO with a refractive index of 1.497 and a thermo-optic coefficient of −4.93×10^−4^ K^−1^ (∂n/∂T) (*33*), at t = 12 μs, the stimulated Raman induced heating causes a refractive index change by −0.07% at the focal center. As shown in Fig. 1D, such an index change nonlinearly extends to the surroundings through heat propagation, thus creating a thermal lens. Formation of such a thermal lens builds the foundation for SRP microscopy.

### Experimental confirmation of the SRP effect

We experimentally confirmed the simulation results with a fluorescence thermometer. It has been well documented that the emission intensities of some fluorophores is temperature dependent (*34*). For instance, the fluorescence intensity of Rhodamine B decreases by ∼2% per Kelvin at room temperature (*35*). This property has been utilized in fluorescence-detected mid-infrared photothermal spectroscopy (*36,37*). Here, we adopt this method to measure the temperature rise at the SRS focus, using Rhodamine B as a fluorescence thermometer. When chirped pump and Stokes lasers are focused into a DMSO solution of Rhodamine B, the Rhodamine B molecules at the SRS focus are electronically excited to emit fluorescence through multiphoton absorption. Meanwhile, when the beating frequency between pump and Stokes is tuned to resonate with the C-H vibration in DMSO, the SRP effect occurs to raise the temperature and accordingly decrease the two-photon fluorescence intensity of Rhodamine B molecules (Fig. 1E).

With this design and identical parameters as in the simulation, we find that the fluorescence intensity drops ∼2.3% after 12 μs of on-resonance SRS, corresponding to ∼1.2 K of temperature rise (Fig. 1F), close to the simulation result of 2.4 K. After weighting each unit in the temperature field with corresponding two-photon excitation intensity of Gaussian beams, the fluorescence change curve can be well fitted as shown in Fig. 1F.

### A chemical microscope sensing the SRP effect

By sensing the local refractive index modulation using a third continuous wave beam, we have built an SRP microscope as illustrated in Fig. 2A. Briefly, the synchronized pump and Stokes pulse trains are intensity-modulated by two AOMs, combined, and chirped to excite the molecules. A probe beam is then collinearly aligned with the SRS beams. A pair of lenses adjusts the collimation of the probe beam to make the probe laser focus axially off the SRS focus, to maximize the photothermal signal (*27*). An iris at the back focal plane of the condenser lens is set to N.A. = 0.4 to convert the probe beam refraction modulation to an intensity modulation.

**Fig. 2.**
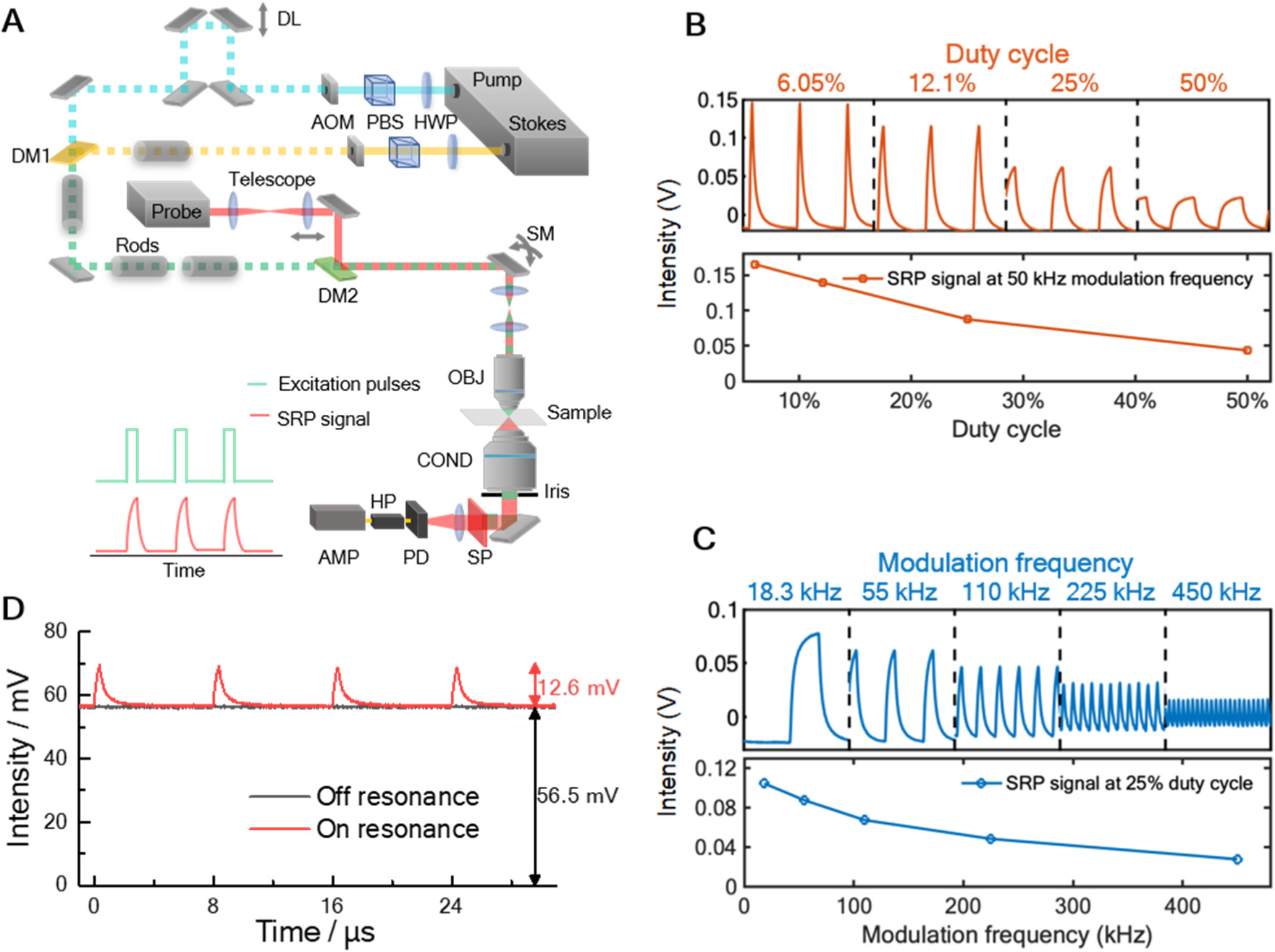
SRP microscope design and characterization of SRP modulation depth as a function of duty cycle and modulation frequency. (**A**) Experimental setup. DM: dichroic mirror; DL: delay line; AOM: acousto-optic modulator; PBS: polarizing beam splitter; HWP: halfwave plate; SM: scanning mirror; OBJ: objective; COND: condenser; SP: spectral filter; PD: photodiode; HP: highpass filter; AMP: amplifier. (**B**) Measured SRP signal as a function of modulation duty cycle. (**C**) Measured SRP signal as a function of modulation frequency. (**D**) SRP generates a large (22.3%) modulation depth with DMSO as the sample. The on-resonance and off-resonance traces were obtained with the Raman shift at 2913 cm−1 and 2850 cm−1, respectively.

The SRP effect creates a divergent lens, which reduces the effective N.A. of the probe laser focus, thus increases the transmission of the probe beam through the iris. A fast photodiode detects the probe beam intensity, followed by a highpass filter and a broadband amplifier. The SRP modulation induced by synchronized pump and Stokes pulses is digitized in real time by a high-speed digitization card. Details can be found in Methods and Fig. S2.

Unlike SRS, both pump and Stokes beam are intensity-modulated in the SRP microscope. Since the SRS intensity is proportional to the product of pump and Stokes peak power, with conserved average laser power, reduction of laser duty cycle leads to higher laser peak power, hence more SRS energy deposition. Our experiment (Fig. 2B) confirmed this idea and showed much higher SRP signal intensity with lower duty cycle. In SRP imaging applications, the duty cycle was set to 5∼10 % as a compromise between signal intensity and laser power. Another key parameter is the modulation frequency. Lower frequency shows higher signal intensity (Fig. 2C) due to longer heat accumulation time, but suffers more from the 1/*f* laser intensity noise (*38*) and meanwhile reduces the imaging speed. 125 kHz was chosen to balance these factors.

For pure liquid, under 5% duty cycle and 125 kHz modulation frequency, the induced modulation on the probe beam was so strong that we could directly measure the SRP signal in the direct current channel without any amplification, as shown in Fig. 2D. With reasonable laser powers on sample to excite the C-H symmetric stretching mode (2913 cm^−1^) of DMSO, the modulation depth reached 22.3%, which is 500-fold higher than the SRS modulation depth (0.04%) with identical average power. The tremendously higher modulation depth builds the foundation for a better detection sensitivity.

In addition to duty cycle and modulation frequency, medium is another crucial factor that affects the photothermal signal intensity (*39*). It has been well-formulated that the photothermal signal intensity (*Σ*_PT_), thermo-optic coefficient (∂*n*/∂*T*) and heat capacity (*C*_p_) hold the relationship of *Σ*_PT_=*n*(∂*n*/∂*T*)/*C*_p_. Yet, the most common medium in biological samples, i.e. water, has a relatively low thermo-optic coefficient (−1.13×10^−4^ K^−1^) (*40*) and relatively high heat capacity (4181 Jꞏkg^−1^ꞏK^−1^). To avoid the signal loss caused by water medium, we investigated a few common liquid media, as shown in table. S1. It is found that glycerol maintains the signal intensity very well. Meanwhile, glycerol shows high bio-compatibility, and is widely used as a mounting medium or clearing agent in bio-imaging. Our simulation of the thermal lens comparison, shown in Fig. S4, also agrees with the previous theory that the peak refractive index change in glycerol medium is ∼2.5-fold as high as in water, with identical heat source (100 nm PMMA nanoparticle under on-resonance SRP heating). Therefore, glycerol was chosen as the medium in following SRP experiments except for the liquid samples. Also, considering the Raman-active vibrational features of glycerol itself, deuterated glycerol (glycerol-d8) was applied for SRP measurement at the C-H and fingerprint regions.

### Limit of detection and spatial resolution

We characterized the performance of our SRP microscope with well-defined samples. We first measured the limit of detection (LOD) for DMSO, focusing on the 2913 cm^−1^ mode. To keep the thermal and optical properties constant throughout the measurement, deuterated DMSO (DMSO-d6) was used as the solvent to dilute DMSO. As shown in Fig. 3A (and Fig. S5 for complete measurement), the SRP spectrum was clean and smooth with high concentration DMSO, and the signal was observable with concentration as low as 15.4 mM. We calculated the LOD with the equation of LOD = 3*σ*/*k*, where *σ* is the standard deviation of the baseline, and *k* is the slope of the intensity-concentration linear calibration curve. Calculation yielded a sub-millimolar level LOD value of 0.93 mM. In comparison, the LOD by SRS under identical average laser powers was found to be 39 mM. Thus, SRP measurement offers a ∼40-fold improvement in LOD. The LODs for C≡C and C-D bonds were measured by using 1,7-Octadiyne in DMSO medium (Fig. S6) and DMSO-d6 in DMSO medium (Fig. S7). In both cases, SRP showed superior sensitivity than SRS with conserved average power.

**Fig.3.**
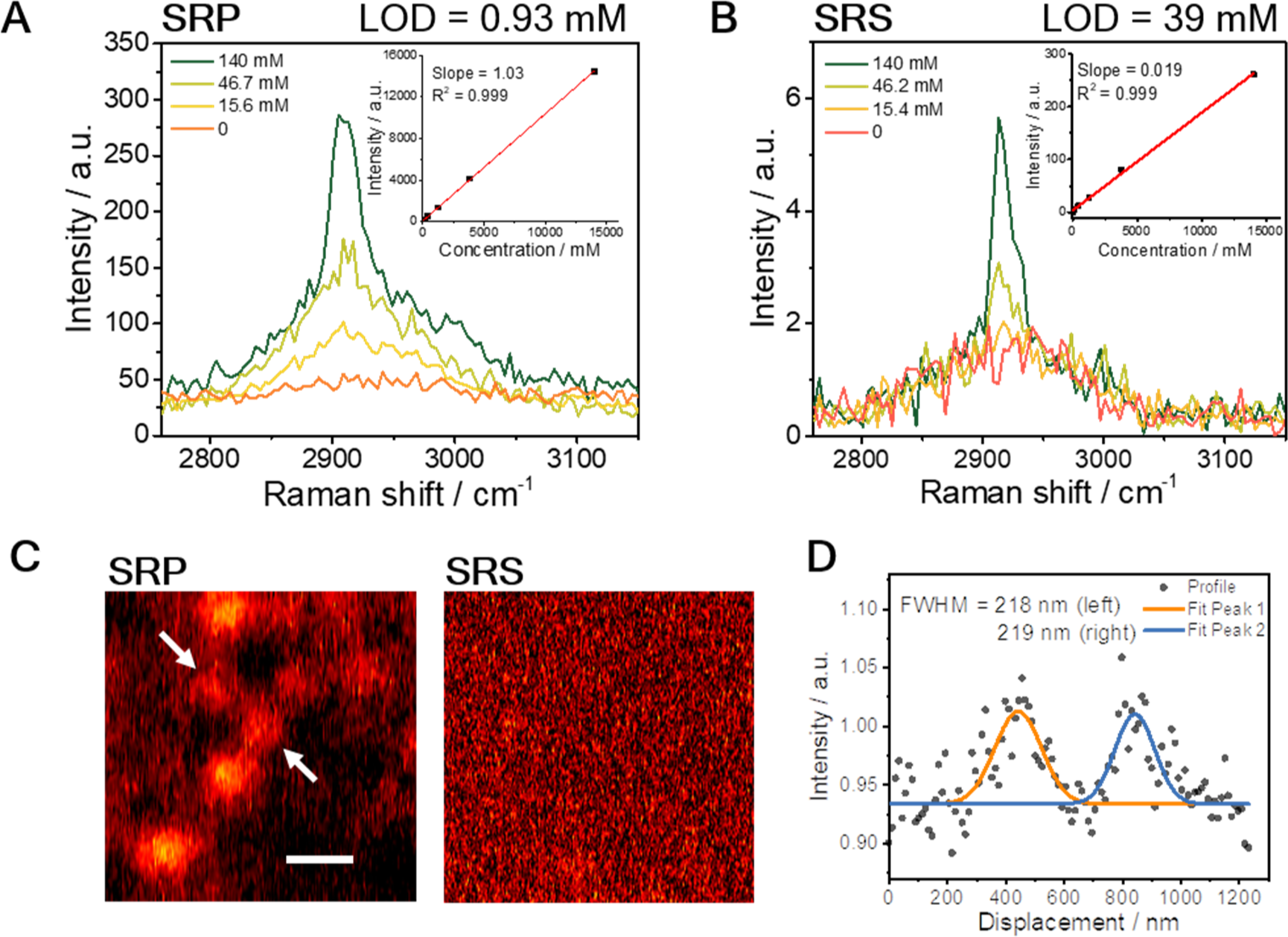
SRP spectroscopy and imaging performance characterization. (A-B) SRP (A) or SRS (**B**) signal with gradient concentration of DMSO dissolved in DMSO-d6. Concentration unit: mM. Insert shows the signal intensity as a function of concentration (complete data in Fig. S6). (**C**) SRP and SRS image of 100 nm PMMA beads at 2930 cm^−1^ with the same average power, at the same field of view. Scale bar: 500 nm. (**D**) The Gaussian fitting measured size of the bead is 218 nm (FWHM).

Such sensitivity improvement allows high-quality imaging of nanoparticles. With the SRP microscope, we successfully acquired hyperspectral image of 100 nm Poly(methyl methacrylate) (PMMA) beads as shown in Fig. 3C. The acquired SRP spectrum showed Raman peak of PMMA at 2950 cm^−1^ which was well-distinguished from the background spectrum as shown in Fig. S8, with an SNR ∼7.0 after BM4D denoising (*41*). In comparison, the SRS measurement showed no contrast of 100 nm beads on the same sample with identical average laser power.

Collectively, SRP showed significantly improved sensitivity comparing to SRS, both for liquid samples and for nanoparticles. Importantly, the introduction of a third probe beam at a shorter wavelength helped improve the spatial resolution (Fig. 3D). We plotted the intensity profile across a pair of 100 nm PMMA beads, with the Gaussian fitted FWHM found to be ∼218 nm. Deconvolution with the beads size generated a FWHM of ∼ 194 nm, which was below the theoretical resolution limit of SRS under the same condition (∼217 nm, FWHM of Airy disk). The FWHM was a little larger than the theoretical resolution limit yielded from the product of pump, Stokes and probe point spread functions (∼167 nm), probably due to the imperfect overlap between the probe and the SRS focus along the axial axis.

### Biological applications

The outstanding performance encouraged us to explore the potential of SRP in bio-imaging. Inspired by the sensitive imaging on 100-nm nanoparticles, we first tested the capability of imaging viral particles. As shown on Fig. 4A1, single varicella-zoster virus (diameter ≈ 180 nm)42 could be clearly resolved from the background with an SNR ∼20. The SRP spectrum of a single virus (Fig. 4A2) peaked at 2950 cm-1, indicating strong contribution from the nucleotide at the core of the virus.

We then applied SRP microscopy to the spectrally silent region. Bacteroides cultured in heavy water were imaged as shown in Fig. 4B1. Decent contrast was obtained from the C-D vibration, while the off-resonance image (Fig. 4B2) showed little signal at the same field of view, confirming the Raman origin of the signals. Such results indicate the potential of SRP in metabolic imaging of single bacteria.

**Fig.4.**
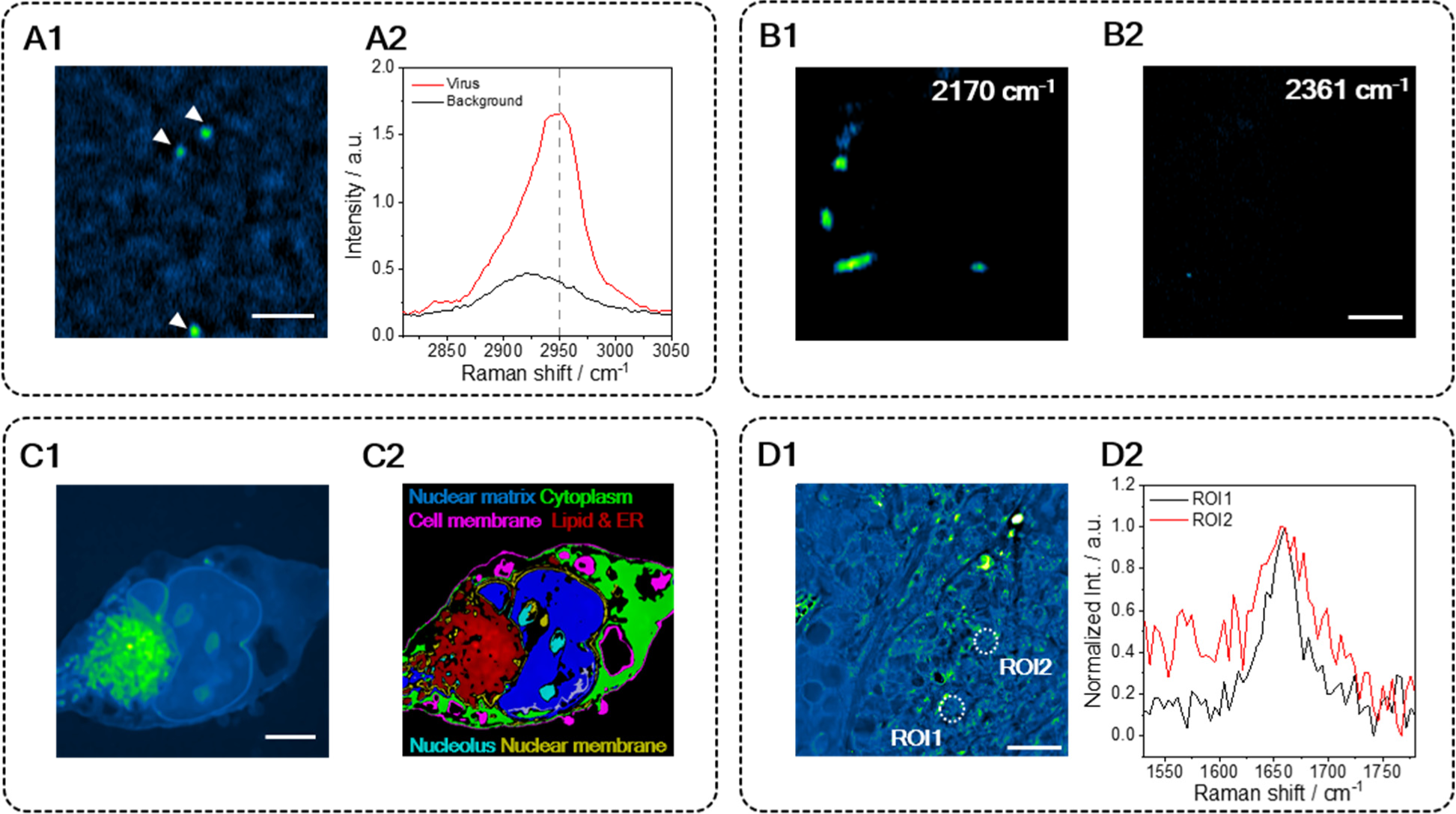
Bio-application of SRP spectroscopic imaging. (**A1**) SRP imaging of a single varicella-zoster virus, 2950^−1^ cm. Scale bar: 2 μm. (**A2**) Single virus Raman spectrum at C-H region, acquired from a single virus in (**A1**). (**B1**) C-D imaging of heavy water cultured Bacteroides at 2170^−1^ cm ((**B1**) On-resonance) or 2361^−1^ cm ((**B2**) Off-resonance). Scale bar: 3 μm. (**C1**) SRP imaging of fixed Mia-Paca2 cell immersed in glycerol-d8, at 2950^−1^ cm. Scale bar: 5 μm. (**C2**) Color coded chemical map through manual phasor segmentation of (**C1**). (**D1**) SRP spectroscopic imaging of OVCAR-5 tissue, cleared with glycerol-d8, at 1650^−1^ cm. (**D2**) Single pixel spectrum at the circled region of interests (ROI).

Pancreatic cancer cell MIA Paca-2 was selected as the testbed for mammalian cell imaging (Fig. 4C1). Glycerol-d8 was applied to replace the PBS buffer and immerse the cells to enhance the SRP contrast. SRP provided nice contrast at the C-H region. Phasor analysis was applied to segment the cellular compartments (Fig. S9), where up to 6 different components could be well identified (Fig. 4C2). Notably, the nuclear membrane outstood from the cytoplasm and the nuclear matrix, giving rise to the potential of applying SRP to study fine structure in a membrane. The high contrast is probably due to the high thermo-optic coefficients and low heat capacities of membranes.

The high sensitivity of SRP also provides access to weak Raman bands in the fingerprint region. Fig. 4D1 showed the SRP image of a 10 μm thick OVCAR-5 cancer tissue at 1650 cm-1, targeting the Amide I band in proteins and the C=C vibration in lipids. High-quality spectra were resolved from the hyperspectral image stack. Lipid (region of interests 1 (ROI1)) and protein (ROI2) species are clearly differentiated, as shown in Fig. 4D2. Collectively, versatile bio-imaging applications are demonstrated, scaling from single virus to cancerous tissues.

## Acknowledgments

We thank Fátima C. Pereira from Centre for Microbiology and Environmental Systems Science University of Vienna for assisting with preparation of bacterial cells.

## Author contributions

J.X.C. conceived the concept of photothermal detection and supervised the research team; Y.Z. built the simulation model, designed and built the SRP imaging system; X.G. implemented the data analysis algorithm; Y.Z. and X.G. performed the imaging experiments and data analysis; J.Y. designed and implemented the matched filtering algorithm; H.N. designed and implemented the broadband detector; H.L. conceived the idea of introducing SHG probe laser; L.W. implemented the SHG probe laser; Y.T. prepared the Mia-Paca 2 cell sample; C.P.D prepared the OVCAR-5 tissue sample. Y.Z., X.G. and J.X.C. wrote the manuscript with input from all authors.

## Data and materials availability

All data and image reconstruction codes are available upon reasonable request to the corresponding author ( (J.X.C.)).

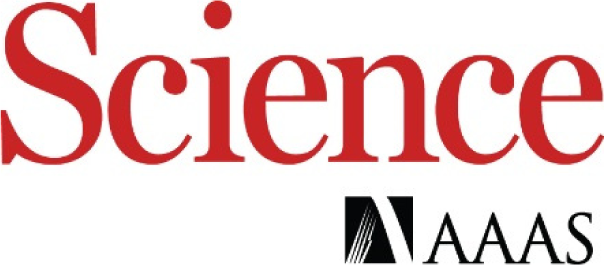

## Supplementary Materials for Photothermally Detected Stimulated Raman Microscopy towards Ultrasensitive Chemical Imaging

### Materials

#### Cell culture and sample preparation

SKOV3 (Cat#: HTB-77) and Mia Paca2 (Cat#: CRL-1420) cells were from the American Type Culture Collection (ATCC). Cells were cultured in high-glucose DMEM medium (Gibco) supplemented with 10% FBS and 100 units/mL penicillin/streptomycin and maintained in a humidified incubator with 5% CO2 supply at 37°C. After overnight seeding in a sterile 35 mm glass-bottom dish (Cellvis) or #1 cover glass (VWR), cells were fixed with 10% neutral buffered formalin for 30 min followed by 3 times PBS wash. Then the cells were covered with glycerol-d8 before sealing and imaging.

#### Tissue sample preparation

A fresh ovarian tumor section was extracted from NU/J mouse (4 weeks old female homozygous for Foxn1nu) purchased from the Jackson Laboratory. The mouse was inoculated with OVCAR5-cisR cells. The sections were immediately fixed in a 10% formalin solution. The extracted tissue section was then washed using PBS solution and cryopreserved by incubation in 15% sucrose solution for 12 hours, followed by incubation in 30% sucrose solution overnight at room temperature. The tissue section was frozen at −80°C by embedding in OCT (Optimal Cutting Temperature) compound in a tissue mold. The tissue section was then sliced using a Microm HM525 cryostat at the Bio-Interface and Technologies Facility, Boston University, into 10 µm thick layers, each placed on separate glass slides and frozen at −80°C until the experiment. To prepare the samples for imaging, the tissue slides were washed using PBS solution to wash off the OCT. The tissue slides were then covered with glycerol-d8 before placing a coverslip to seal the glycerol-d8 covered tissue layer.

#### Glycerol-agar (glycerol-d8-agar) medium preparation

1% (w/w) agar (Sigma Aldrich) was mixed with glycerol (or glycerol-d8, Sigma Aldrich), then microwaved for 2min to fully dissolve the agar.

#### Nanoparticle, bacteria, and virus preparation

For 100 nm PMMA nanoparticles, 10 μL solution was mixed with warm glycerol-d8-agar medium, then sandwiched between two No.1 coverslips before imaging. For D_2_O cultured bacteroid (*15*), 10 μL solution of fixed bacteria was mixed with warm glycerol-agar medium, then sandwiched between two No.1 coverslips before imaging. Varicella zoster virus solution (Fisher Scientific) was dropped onto a No.1 coverslip and dried on top. The sample was covered with glycerol-d8, then sandwiched between two coverslips and sealed with nail polish before imaging.

### Methods

#### Modeling the temperature change induced by SRS

To quantitatively evaluate the SRP effect, we built a model to simulate the heat deposition spatially and temporally. The SRS induced temperature change is dependent on the pulse width, pulse energy, laser repetition rate, and thermal properties (thermal conductivity, heat capacity, etc.) of the sample and the surrounding medium. First, the deposited energy by SRS is estimated by the modulation depth on the pump or Stokes beam along with the pulse energy. Then, to simulate the thermal conduction, the time domain and spatial domain are divided into finite elements for the calculation according to Fourier’s law: *dQ* = —*kA* (*dT*⁄*dr*), where *dQ* is the conducted heat energy in the time window, *k* is the thermal conductivity, *A* is the surface area, *dT* is the temperature difference between distance *dr*. The SRS on-off process induces a heating and cooling process at the focus. Heat transfer happens during both heating and cooling. During the heating process, the SRS pulses deposit heat to the sample instantaneously with a Gaussian distribution as the vibrational excited state relaxation time is much faster compared to the simulation time grid. Finally, with the heating estimated by the modulation depth and the thermal conduction calculated according to Fourier’s Law, the temperature spatial distribution at each time point is calculated.

To simplify the model, an isotropic Gaussian SRS heating area is assumed. The simulation area is set to be a 48-layer model with increasing step size from the center to the edge (Fig. S1). The time step is set to be 200 ps, which is much smaller than the heat propagation time to ensure a converged result. DMSO has a heat conductivity of 0.2 W/(mꞏK). The laser parameters are set to 50% duty cycle, 80 MHz repetition rate, according to the commonly used SRS setting.

To predict the fluorescence change upon temperature rise, the excitation profile of two photon fluorescence was modeled as the normalized Gaussian point spread function of the objective. Weights based on the assumed excitation profile were put onto the temperature rise of each layer, then multiplied by the −2%/K rate (*34*), to yield the fluorescence change.

#### Measurement of temperature at the SRS focus

A fluorescence thermometer, Rhodamine B (−2%/K) (*34*), is introduced to measure the temperature rise at the SRS focus. 80 μM Rhodamine B dissolved in DMSO is sandwiched between two thin coverslips (No.1; Thermo Fisher) for the measurement. A dual-output synchronized laser source (InSight X3; Spectra-Physics) provides pump and Stokes beams, respectively. The Stokes beam is modulated by an acousto-optic modulator (1205c; Isomet Corporation) at 40 kHz with the first order beam to provide a 100% modulation depth. Then the pump and Stokes beams are chirped by 75 and 90 cm glass rods (SF57; Scott AG), respectively, to implement hyperspectral SRS under a spectral focusing scheme. The path length of the Stokes beams could be adjusted by a motorized delay line (X-LRM025A-KX13A, Zaber Technologies). The two beams are combined by a dichroic mirror (950 nm; Chroma) and then collinearly guided into a laser scanning microscope. To vibrationally excite the C-H bond in DMSO, the pump laser was set to 800 nm with the Stokes beam wavelength fixed at 1045 nm. These two beams could also simultaneously excite the two-photon fluorescence signal of Rhodamine B. A 40 water objective (numerical aperture (N.A.) = 0.8; Olympus) focused the two beams onto the sample, with power of 25 mW for pump and 50 mW for Stokes. The output light is collected in the forward direction by an oil condenser (N.A. = 1.4, Aplanat Achromat 1.4; Olympus) and a 75 mm A-coating focal lens (Thorlabs). A silicon photomultiplier (C14455-3050GA; Hamamatsu) module with a bandpass optical filter (RT570/20x; Chroma) and two short-pass filters (1000SP, 775SP; Thorlabs) is used to detect the fluorescence signal. The output of the silicon photomultiplier is recorded by a spectrometer (Moku:Lab, Liquid Instruments) for the analysis in Fig. 1F.

#### SRP microscope

The SRP microscope is based on the SRS setup described in the previous section. Both pump and Stokes beams are modulated by two synchronized acousto-optic modulators outputting the first order beams, various duty cycles, and modulation frequency at 125 kHz. Besides, a 765 nm continuous probe laser (TLB6712-D; Spectral Physics) is added after the combining dichroic mirror by a polarized beam splitter to form a three beam copropagating colinear system (Fig. 2B). The three beams are guided to a two-dimensional galvo scanning unit (GVS002; Thorlabs), which is conjugated by a four-focal system to the back aperture of a 100× oil objective (N.A. = 1.49, UAPON100XOTIRF; Olympus). The N.A. of the condenser is adjusted to 0.4 to enable the detection of the thermal lensing signal. The detector is a broadband silicon photodiode (Hamamatsu) with 50-ohm resistance, a 22 kHz highpass radio frequency filer (Mini-circuits) and a 46 dB low noise amplifier (SA230-F5; Wayne). A tilted bandpass optical filter (FL780-10; Thorlabs) is mounted before on the detector to block the SRS beams and allow sole detection of the probe beam.

The output signal is digitized by a fast data acquisition card (Alazar card, ATS9462; Alazar Technologies). In imaging the virus on a coverslip, the 1045-nm femtosecond laser output is sent to an LBO crystal to generate 522.5 nm SHG laser beam as the SRP probe. This femtosecond probe with reduced coherence length minimizes the interference from the medium/coverglass interface.

When performing SRS imaging on the same setup, the probe laser is turned off. A 1.4 N.A. condenser is used to minimize the cross phase modulation background. The modulation on the pump is set to always on and the Stokes was modulated at around 2.25 MHz. After the condenser, two short-pass optical filters (980SP, Thorlabs) pass the pump beam to the photodiode. A laboratory-built resonant amplifier after the photodiode picks up the stimulated Raman loss signal. The photodiode output is sent to the lock-in amplifier (MFLI; Zurich) and the stimulated Raman loss signal is recorded with a data acquisition card (NI DAQ card, PCI-6363; National Instruments).

The SRP system is electronically synchronized by a NI DAQ card. The NI DAQ card controls the galvo scanning unit and generates the transistor-transistor logic trigger to control the sampling of the Alazar card and the pixel trigger to control the function generator. The function generator generates rectangular waves with various duty cycles at 125 kHz in burst mode to control the two acousto-optic modulators to modulate the pump and Stokes pulse trains (Fig. S2). The amplified signal from the detector set is digitized by the Alazar card at a sampling frequency of 20 MHz and then sent to the host computer for further analysis.

#### SRP signal digitization and processing

After recording the signal from each pixel, a Whittaker smoother removes the fluctuated baseline caused by variation in the transmission (*43*). Matched filtering is then applied to improve the SNR (*44*). Fourier transform on the baseline removed signal is applied to generate the signal spectrum. To reduce the out-of-frequency noise, a match multi-bandpass filter at up to seven harmonic frequencies of the modulation frequency is applied to the signal spectrum (Fig. S3). The width of the single bandpass filter is set to 13.89 kHz when the pixel dwell time is 72 μs.

The match-filtered spectrum is then inversely Fourier transformed back to the time domain. The SPR signal intensity of each pixel is calculated from the average peak-dip contrast of the processed time domain data.

#### SRP imaging parameters

The modulation depth is measured with 5% duty cycle modulation, 50 mW and 20 mW for Stokes and pump, respectively. The virus image is acquired with 5% duty cycle modulation, 40 mW and 20 mW for Stokes and pump, respectively. The bacteria image is acquired with 10% duty cycle modulation, 30 mW and 20 MW for Stokes and pump, respectively. The cell image is acquired with 10% duty cycle modulation, 30 mW and 15 mW for Stokes and pump, respectively. The tissue image is acquired with 5% duty cycle modulation, 30 mW and 10 mW for Stokes and pump, respectively. All powers are measured before the microscope.

#### Phasor analysis

Phasor analysis was performed with the standardized phasor analysis plug-in in ImageJ (1.49v). The phasor domain segmentation was performed manually to maximize the separation of different components as well as the integrity of each component.

---

**Fig. S1.**
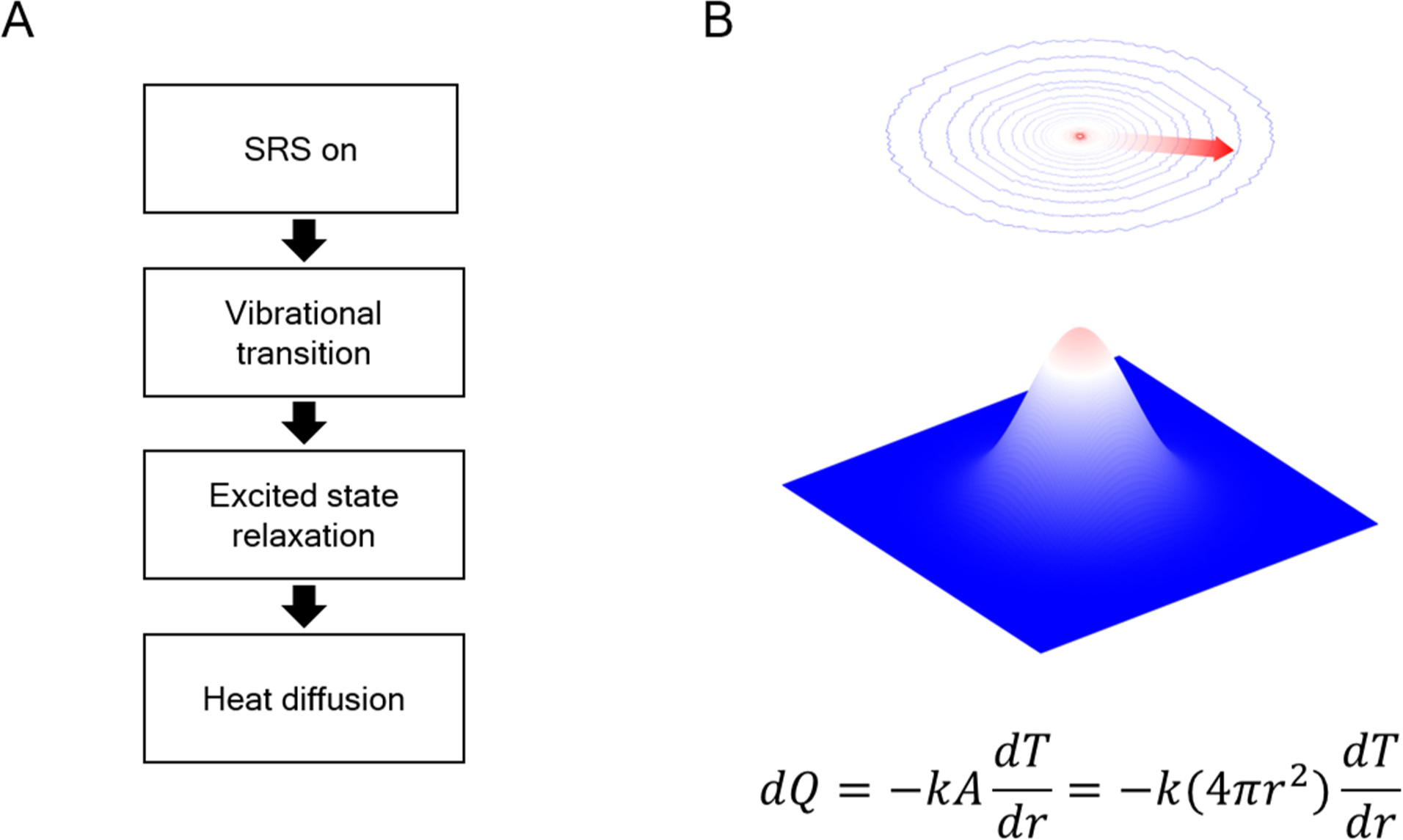
SRP simulation model. The system is gridded into concentric spherical shells. The red arrow indicates the thermal propagation direction.

**Fig. S2.**
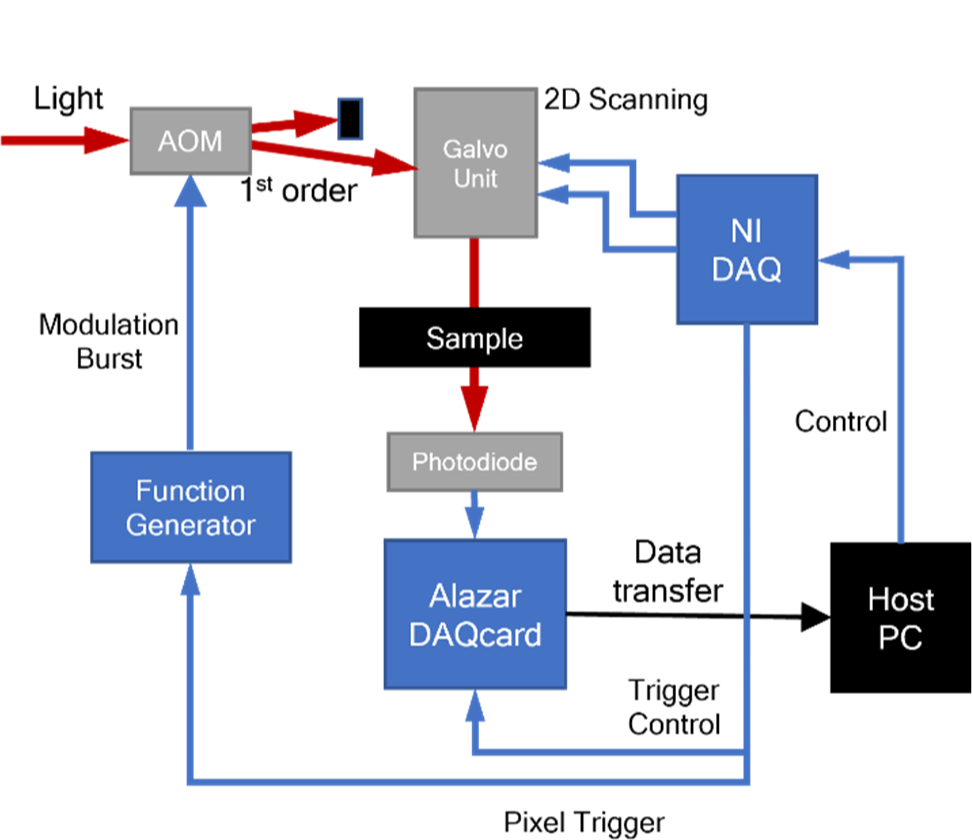
Electronic control and data acquisition flow in the SRP microscope.

**Fig. S3.**
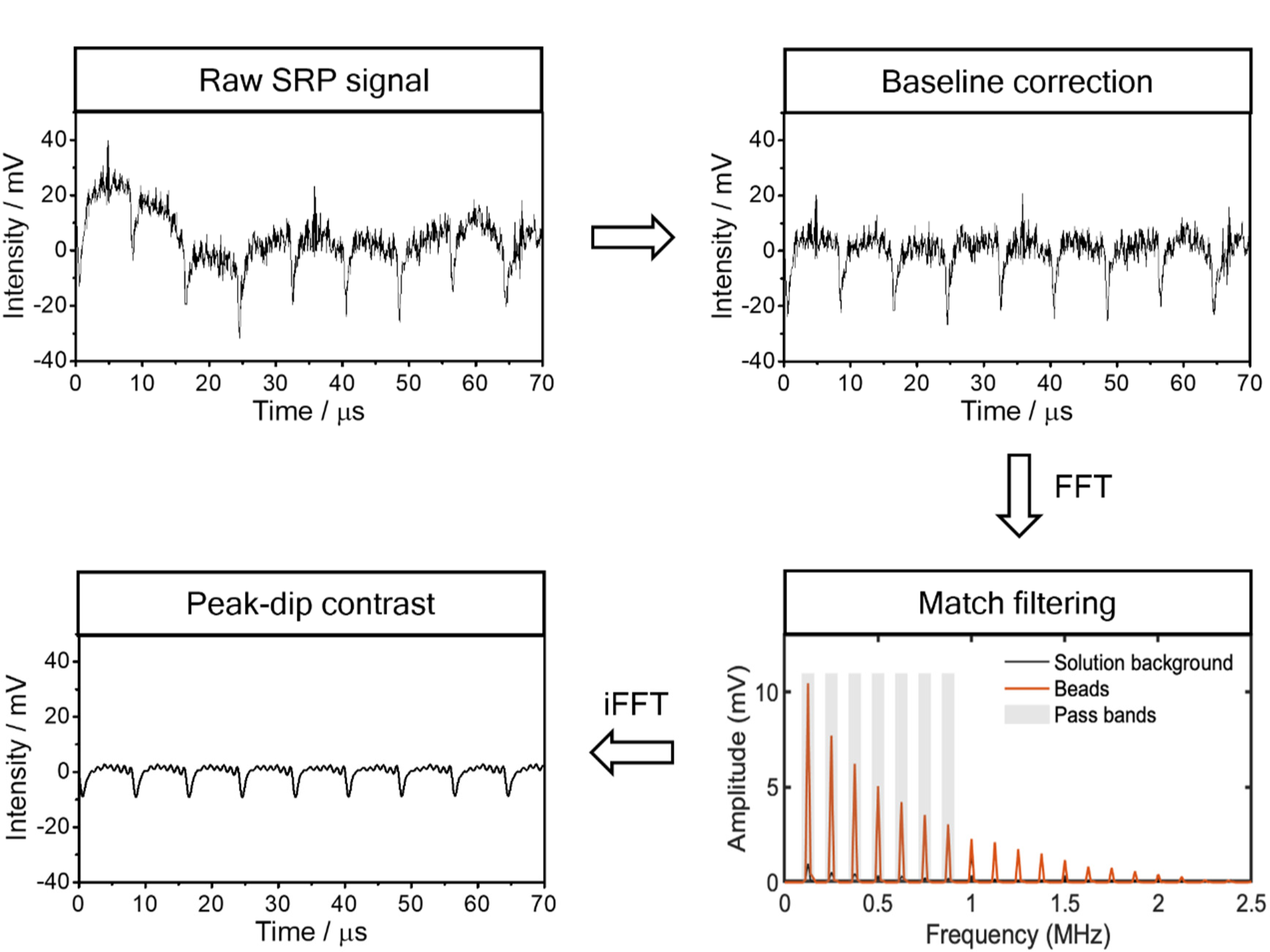
A thermal dynamic trace from SRP measurement and under different stages of data processing. The signal is acquired from a single 100 nm PMMA nanoparticle at 2950 cm-1 immersed in glycerol-d8. FFT: fast Fourier transform; iFFT: inverse fast Fourier transform.

**Fig. S4.**
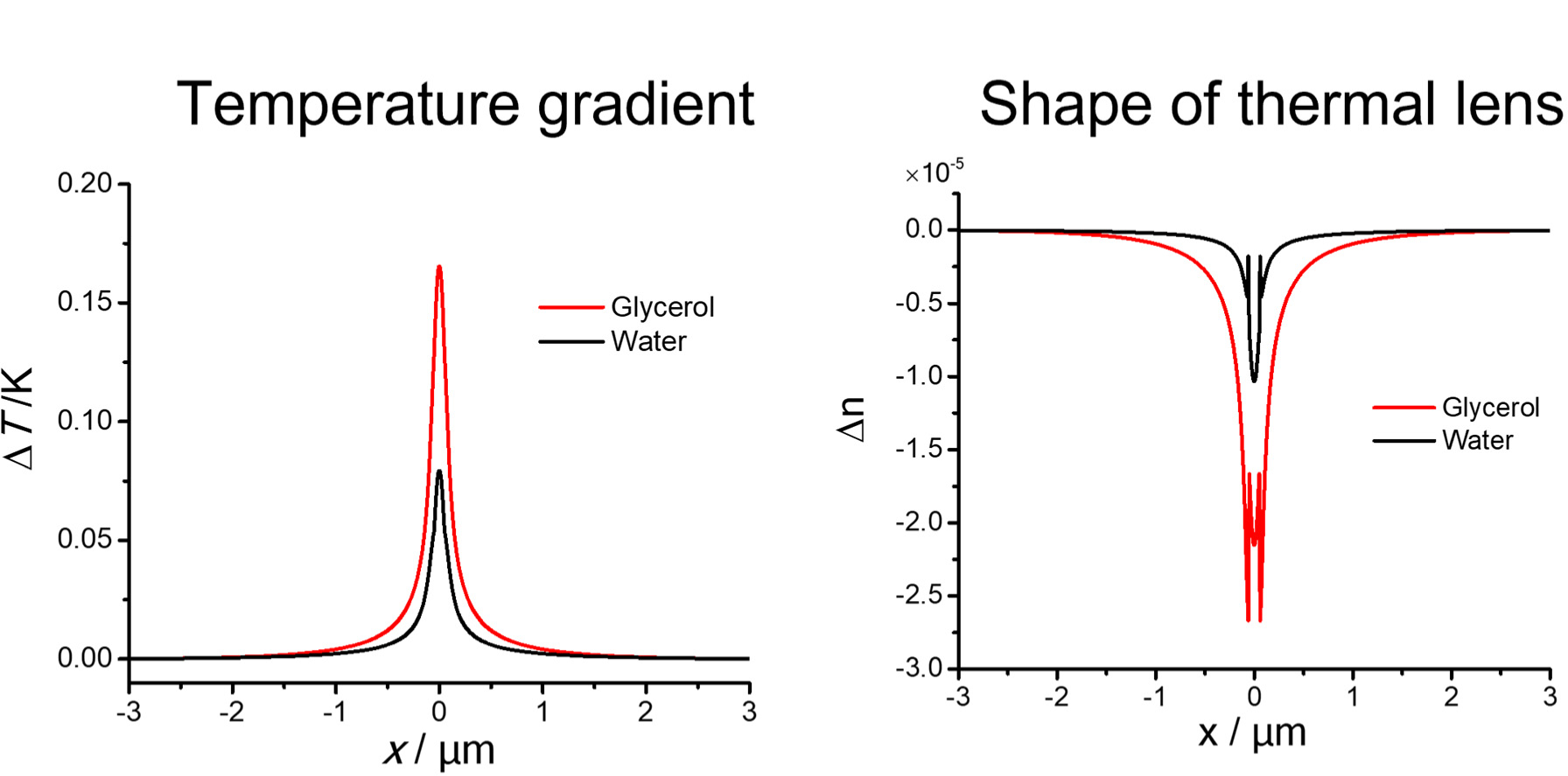
Simulation results for the shape of temperature gradient and thermal lens induced by a 100 nm PMMA particle in glycerol or water.

**Fig. S5.**
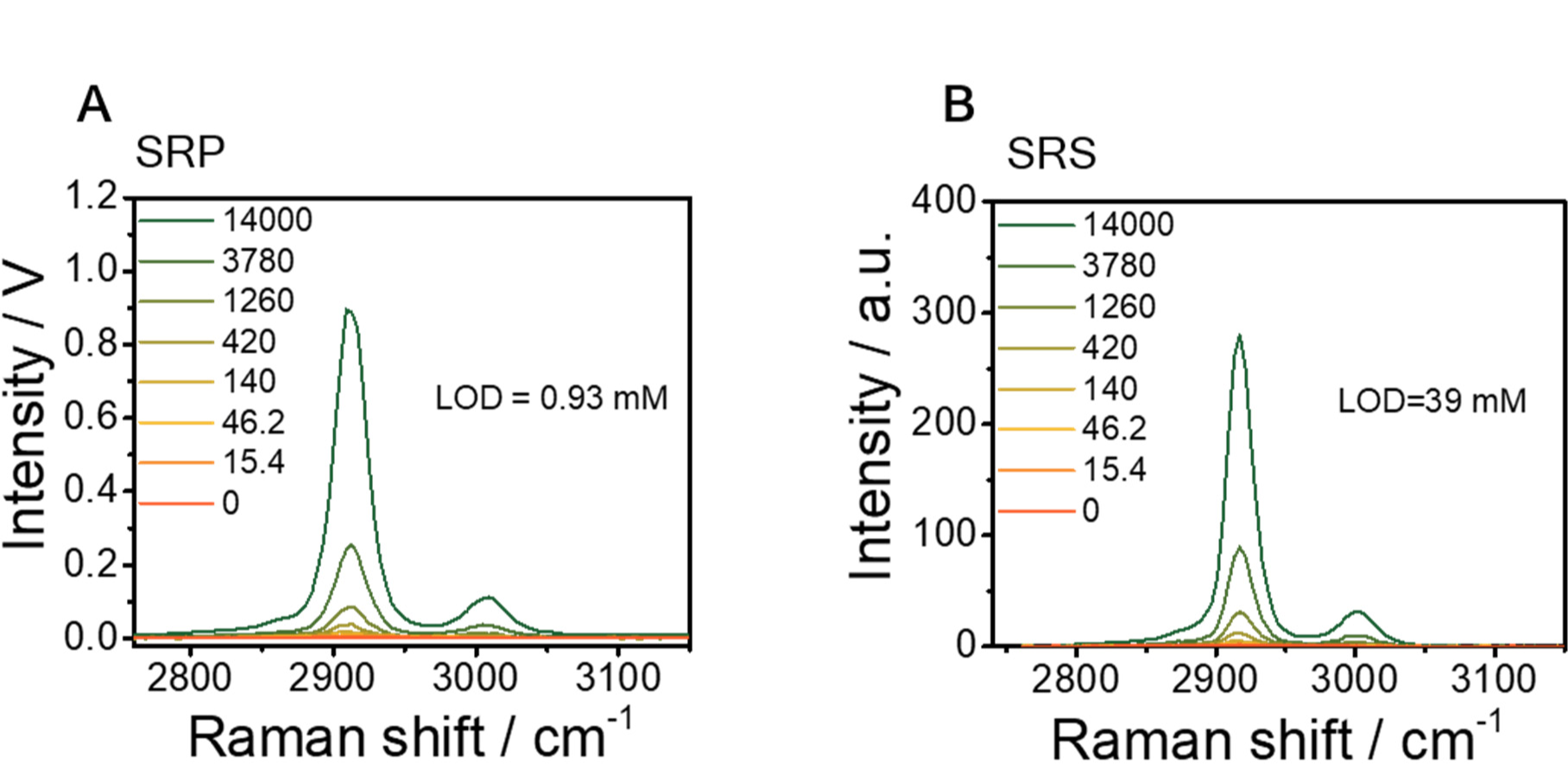
Complete data set of LOD measurement of DMSO dissolved in DMSO-d6 with SRP (A) and SRS (B). Unit of concentration: mM.

**Fig. S6.**
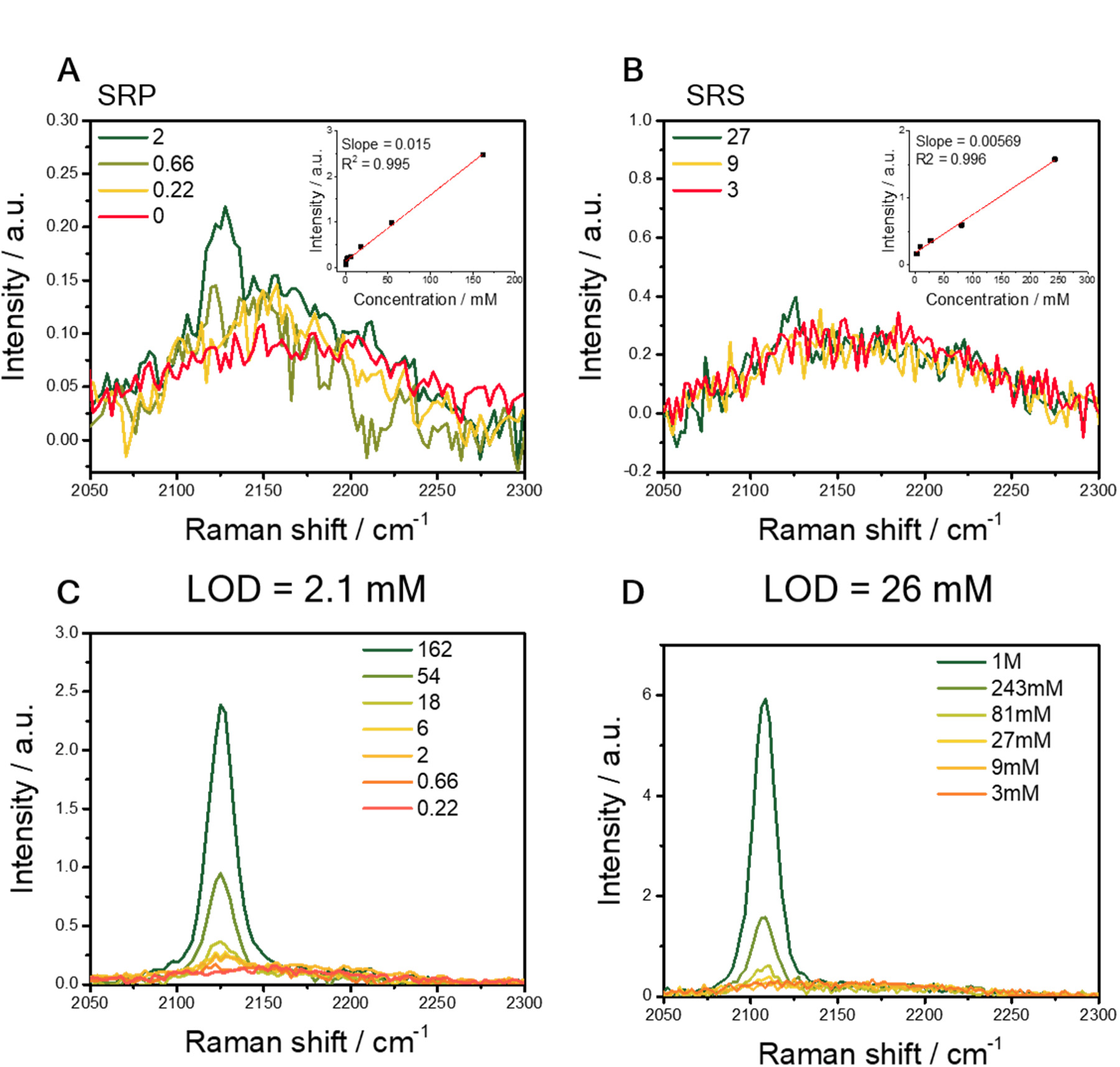
LOD of 1,7-Octadiyne in SRP and SRS measurement. (**A**-**B**). SRP (**A**) or SRS (**B**) signal with a gradient concentration of 1,7-Octadiyne dissolved in DMSO. Concentration unit: mM. Insert shows the signal intensity as a function of concentration. (**C**-**D**). The complete data set of LOD measurement with SRP (**C**) and SRS (**D**). Unit of concentration: mM.

**Fig.S7.**
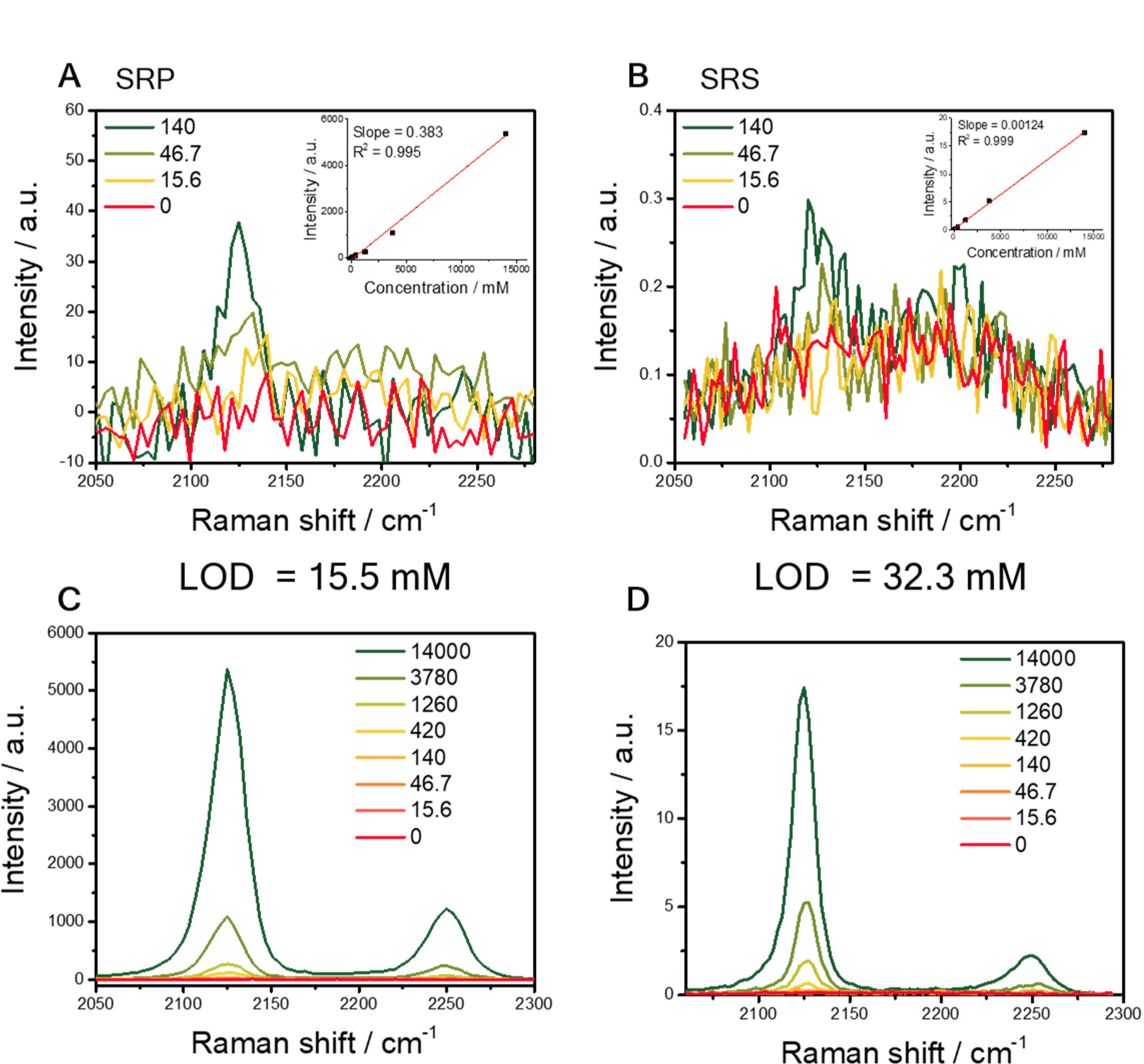
LOD of DMSO-d6 in SRP and SRS measurement. (**A**-**B**). SRP (**A**) or SRS (**B**) signal with a gradient concentration of DMSO-d6 dissolved in DMSO. Concentration unit: mM. Insert shows the signal intensity as a function of concentration. (**C**-**D**). The complete data set of LOD measurement with SRP (**C**) and SRS (**D**).

**Fig. S8.**
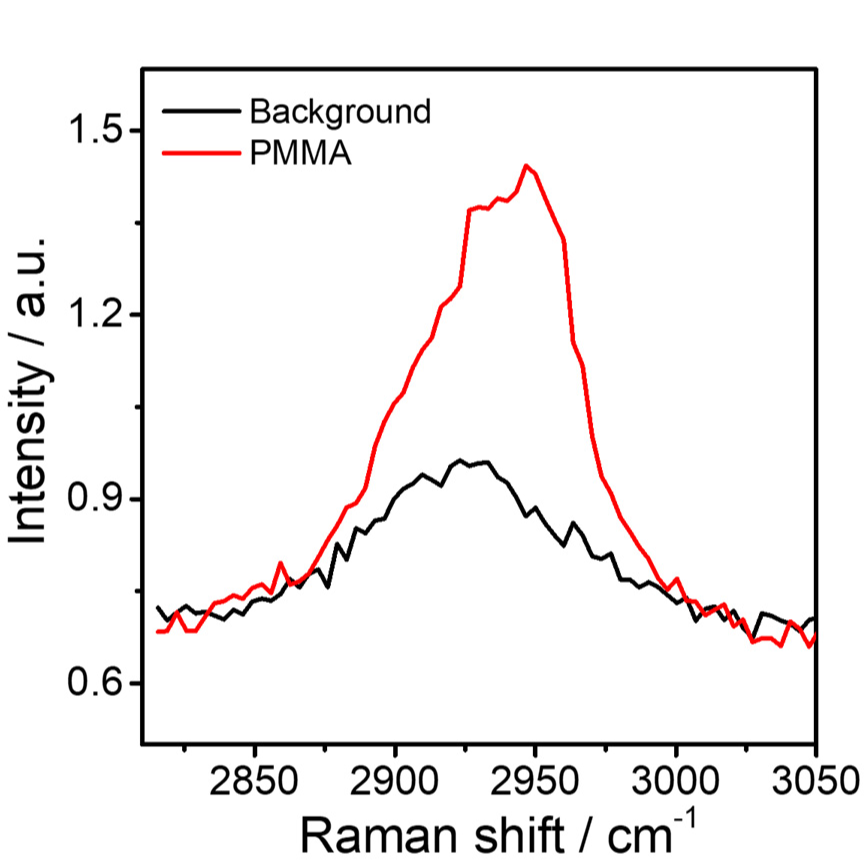
SRP spectrum of a single 100 nm PMMA nanoparticle at the C-H region.

**Fig. S9.**
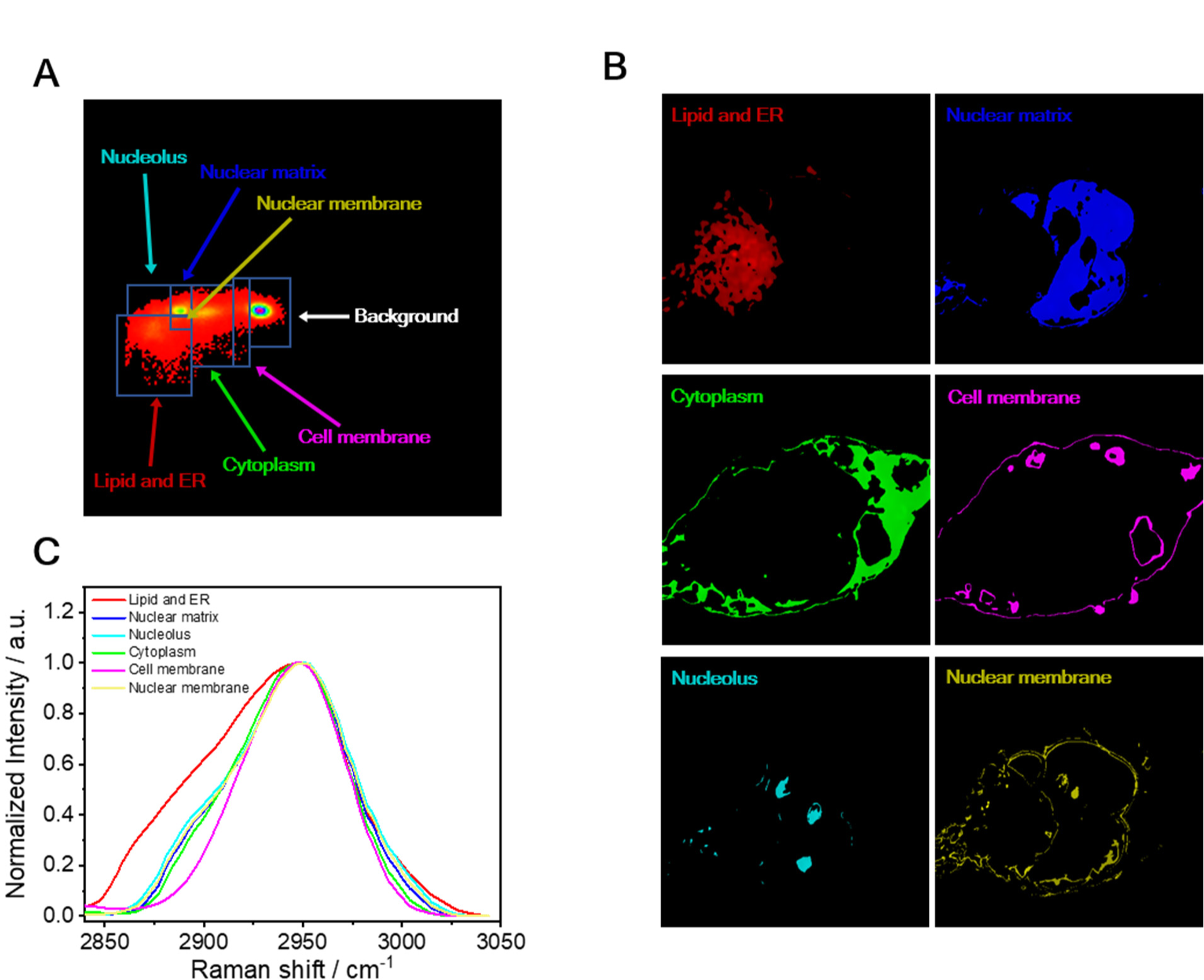
Phasor analysis for the SRP image of glycerol-d8 immersed Mia PACA-2 cells. (**A**). Segmentation of the SRP image in the phasor domain. (**B**). Mapping of each component from phasor analysis. (**C**). The SRP spectrum at C-H region for each component.

**Table S1.** Thermal and thermo-optic properties and estimated relative signal intensity of the mediums related to the study.

| **Thermal property** | **Unit** | **DMSO** | **hexane** | **glycerol** | **water** |
| --- | --- | --- | --- | --- | --- |
| **Heat Capacity** | **J/(kg·K)** | **1966** | **2251** | **2400** | **4184** |
| **Thermal conductivity** | **W/(m·K)** | **0.200** | **0.124** | **0.283** | **0.598** |
| **Thermo-optic coefficient**<br>**dn/dT (10<sup>-4</sup>)** | **K<sup>-1</sup>** | **-4.93 (33)** | **-5.20 (43)** | **-2.30 (44)** | **-1.13 (40)** |
| **Refractive index** |  | **1.479** | **1.375** | **1.473** | **1.333** |
| **Relative signal intensity** | **a.u.** | **10.3** | **8.83** | **4.27** | **1** |

